# Increased temperatures reduce the vectorial capacity of *Aedes* mosquitoes for Zika virus

**DOI:** 10.1101/702431

**Authors:** Onyango Maria Gorreti, Bialosuknia Sean, Payne Ann, Mathias Nicholas, Kuo Lilli, Vigneron Aurelien, DeGennaro Matthew, Ciota Alexander T, Kramer Laura D

## Abstract

Rapid and significant range expansion of both ZIKV and its *Aedes* host species has resulted in ZIKV being declared a global health threat. Mean temperatures are projected to increase globally, likely resulting in alterations of the transmission potential of mosquito-borne pathogens. The relationship between temperature and ZIKV transmission has not been well characterised for *Aedes aegypti* and *Aedes albopictus*.

To understand the effect of diurnal temperature range on the vectorial capacity of *Aedes aegypti* and *Aedes albopictus* for ZIKV, factors contributing to transmission potential were measured at different temperature regimens. Their longevity and blood feeding rates were assessed, and vector competence was determined following feeding on blood meals with 8.3 log_10_ PFU/ml ZIKV.

Higher temperature resulted in decreased longevity of *Ae. aegypti* [Log-rank (Mantel-Cox) Test, Chi-square, df 35.66, 5 P (<0.0001)] and a significant decrease in blood feeding rates across groups [Z score (−5.8478) P (0.0444)]. Temperature had a population and species-specific impact on ZIKV infection rates. Overall, *Ae. albopictus* reared at the lowest temperature regimen demonstrated the highest vectorial capacity (1.63) and the highest transmission efficiency (57%). Temperature increases decreased vectorial capacity across groups, yet the largest decreases were measured for *Ae. aegypti*.

The results of this study suggest that future climate change could significantly impact vector competence, blood feeding behavior and longevity, and therefore decrease the overall vectorial capacity of *Aedes* mosquitoes. It is also clear that this impact is likely to be both species and population-specific.

## Introduction

Zika virus (ZIKV; *Flavivirus, Flaviviridae*), which prior to 2007 was geographically limited to Africa and Asia (1–3) has undergone an unprecedented range expansion in recent years and evolved into a global health threat. ZIKV was first identified in Brazil in May 2015 and subsequently spread throughout the Americas (3–8). The explosive spread of ZIKV from Asia to the Americas and the association with disease outcomes such as microcephaly among infants and Guillain-Barre syndrome among adults necessitates the need to better understand the transmission dynamics of the virus and the potential impact of climate change.

A recent study by Liu et al., 2017 showed that a mutation of the non-structural protein 1 (NS1) may have enhanced the infectivity of ZIKV and facilitated transmission by *Aedes aegypti* in the Americas. Multiple studies have also demonstrated that the geographic origin of both the mosquito and the viral strain affect the vector competence of *Aedes* mosquitoes for ZIKV (1,3,8–10).

Despite progress made to understand the biology of ZIKV infection and its interactions with its vectors, the intrinsic and extrinsic factors that govern the vectorial capacity for ZIKV are still not well understood. Included in this is environmental temperature, which may influence mosquito physiology and viral replication (11) and subsequently have significant effects on virus transmission (12).

It is expected that global climate change will lead to a geographical expansion of tropical disease (particularly vector borne pathogens) throughout temperate regions(13). Higher temperatures are known to speed up biochemical reactions that use up energy, resulting in increases in activity, development and reproduction, yet heightened metabolism can come at a cost. While temperature increases might accelerate viral replication, extrinsic incubation rates and vector factors noted above, there could also be a negative impact on fitness of the vector and its capacity to transmit pathogens (14–17). This study aimed to understand the effect of diurnal, fluctuating temperature regimes on vectorial capacity of *Ae. aegypti* and *Ae. albopictus* for ZIKV. We modelled a 2°C temperature increase, the estimated global increase over the next 50 years(18). *Ae. aegypti* were reared and maintained at high (H) (day 32°C/ night 28°C [D32N28]) and moderate (M) (day 30°C/ night 26°C[D30N26]) temperatures; *Ae. albopictus* at moderate (day 30°C/ night 26°C[D30N26]) and low (L) (day 28°C/ night 24°C [D28N24]) temperature regimes. One population of *Ae. albopictus* (*Ae. albopictus* Long Island, ALB LI) and two populations of *Ae. aegypti* (*Ae. aegypti* Mexico, AEG MX and *Ae. aegypti* Miami, AEG MI) were used in this study and the effect on life-history traits and vector competence were measured. Insights into the impact of future climatic changes on ZIKV transmission dynamics of *Ae. aegypti* and *Ae. albopictus* will improve the epidemiology of this disease.

## Materials and methods

### Mosquitoes

Three populations and two species of *Aedes* mosquitoes originally collected from three distinct geographical and environmental regions (Figure 1) were utilized in this study. *Ae. albopictus* (ALB LI; kindly provided by Illia Rochlin, Suffolk County Health Department) were originally collected in Suffolk County, NY in 2014 and subsequently colonized in the New York State Department of Health (NYSDOH) Arbovirus Laboratory. Mexican *Ae. aegypti* (AEG MX; kindly provided by GD Ebel, Colorado State University) were originally collected in Poza Rica, Mexico, in 2016. *Ae. aegypti* (Miami, AEG MI) were originally collected in Miami-Dade County, Florida in October 2017 (kindly provided by M DeGennaro, Florida International University).

**Figure 1.**
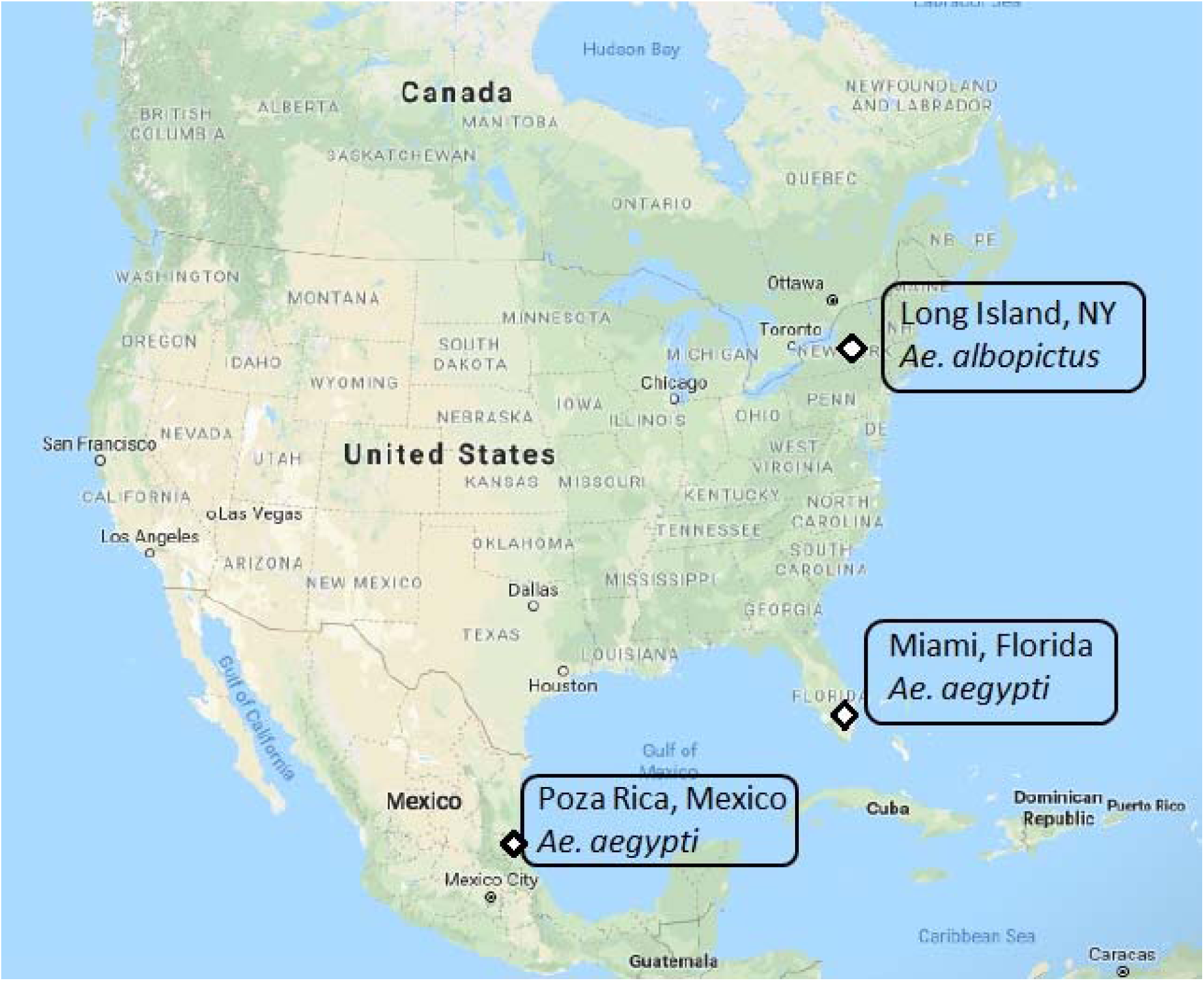
Collection sites of the *Aedes* mosquitoes. The AEG MI mosquitoes were collected from Miami, Florida, AEG MX population from Poza Rica, Mexico while the ALB LI from Long Island New York.The different sites of collection have different climatic conditions that include hot and humid summers, short warm winters in Miami, tropical wet and dry in Mexico and cold and temperate in Long Island, New York. The temperature used to determine the temperature regimes utilised in this study are the estimated temperatures during the peak transmission season. These include; day 30°C, night 26°C in Miami and Mexico while that of Long Island is day 28°C, night 24°C. Source, Google maps.

Colonies were maintained at 27°C under standard rearing conditions before the eggs were hatched. F23 AEG MX, F3 AEG MI and F15 ALB LI eggs were hatched under vacuum pressure in 1 L dechlorinated water initially incubated for 3 h at the three distinct diurnal temperature regimes (Figure 2). Baseline temperature regimes (L for *Ae. albopictus* and M for *Ae. aegypti*) were estimates of mean day and nighttime temperatures during peak transmission in regions from which populations were derived. A subgroup of each population was reared and held at day and night temperatures 2°C higher than baseline (M for *Ae. albopictus* and H for *Ae. aegypti*; Figure 2).

**Figure 2.**
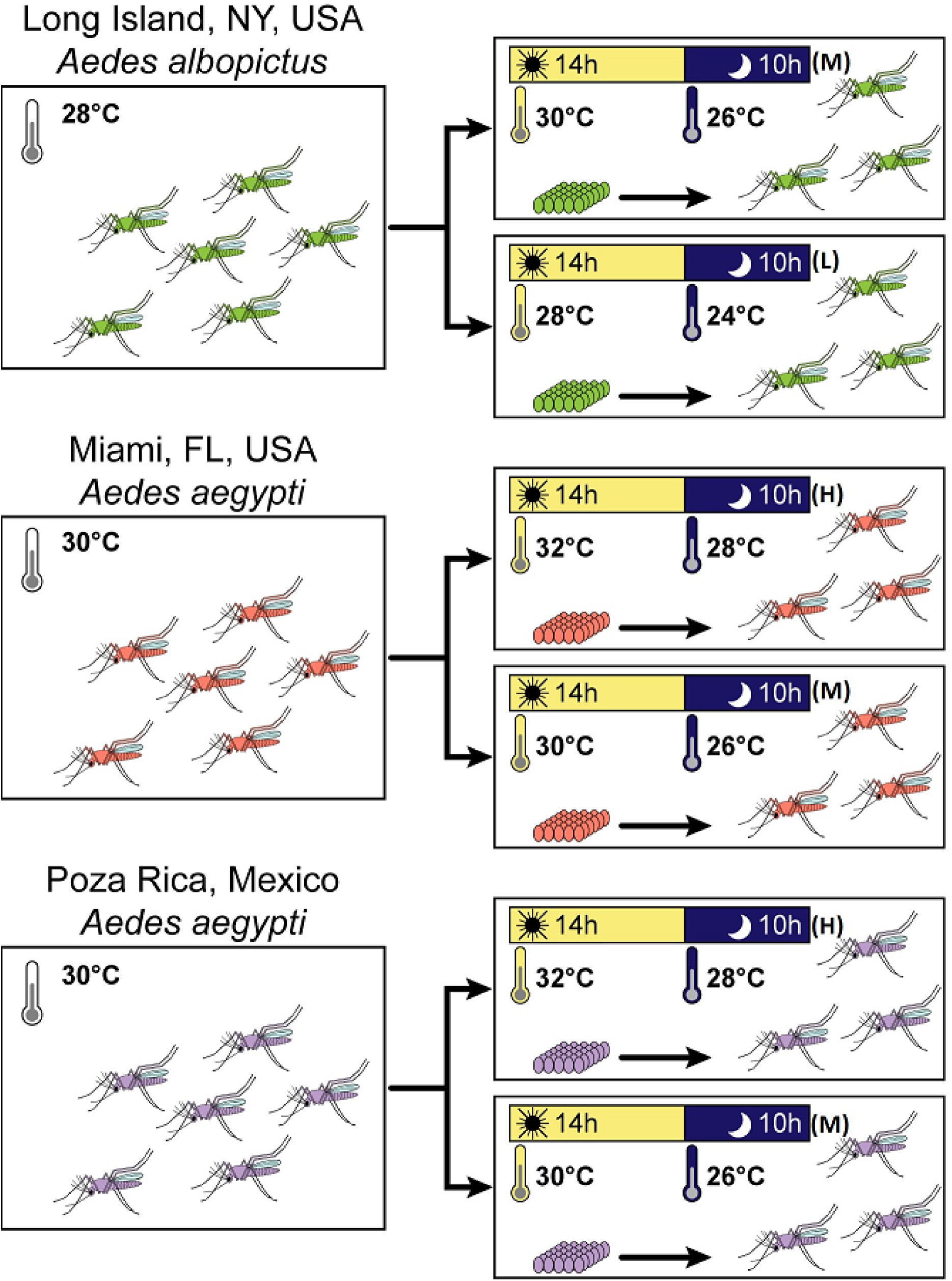
Schematic illustrating the experimental workflow. The temperature regimen selected for each region was based on a 2 °C higher than the day and the night temperature at peak transmission season of each geographical sample origin [D30N26 (M); D32N28 (H) and D28N24 (L)]. The eggs were vacuum hatched after warming the water to the day time temperature and the immature and adult stages reared at the stated temperature regimen.

Collected larvae were maintained in plastic rectangular flat containers [35.6 cm length x 27.9 cm width x 8.3 cm height (Sterilite, catalog no. 1963)] at a density of 200 larvae per 1L of dechlorinated water and reared at 40-60 % relative humidity and a light dark (LD) photoperiod 14:10 h. The larvae were fed 0.25g of Tetra pond Koi growth feed for 1^st^ and 2^nd^ instar larvae and 0.5g for 3^rd^ and 4^th^ instar larvae (10).

Both male and female adults were transferred to 3.8 L cardboard cartons upon emergence and housed together for 9 days under the different temperature regimens (Figure 2) while being provided with sugar and water *ad libitum*. To stimulate blood feeding, the 9-day old females were starved of water and sugar 24 h before infectious blood meal.

### Zika virus vector competence

To test for the effect of temperature, population and mosquito species on vector competence for ZIKV, female *Aedes* mosquitoes were orally exposed to previously frozen 8.3 log_10_ PFU/ml ZIKV HND strain. Zika Virus HND (2016-19563, GenBank accession no. KX906952) was isolated as described in Ciota et al., 2017.

The virus was diluted 1:1 with defribinated sheep blood plus 2.5% sucrose and sodium bicarbonate was included to adjust pH to 8.0. The female mosquitoes were offered the infectious blood meal through a 37°C pre-heated Hemotek membrane feeding system (Discovery Workshops, Acrington, UK) with a porcine sausage casing membrane. After an hour, the mosquitoes were anaesthetized with CO_2_ and immobilized on a pre-chilled tray connected to 100% CO_2._ Engorged females were separated and placed in three separate 0.6 L cardboard cartons (30 individuals per carton). In addition, 1 ml of each blood meal was transferred to a 1.5 ml Safe Seal microtube (Eppendorf, Hamburg Germany) and stored at −80°C to allow for the determination of ZIKV titers. The engorged females were maintained on 10% sucrose solution provided ad *libitum*. The 0.6 L cardboard cartons were kept at the respective temperature regimens (Figure 2).

### Infection and transmission analysis

On day 4, 7 and 14 post infectious blood meal, mosquitoes were immobilized by exposure to triethylamine (Sigma Aldrich, St. Louis, MO, USA), the legs were removed from 30 mosquitoes of each population and species, and placed in 500 µl mosquito diluent (MD; 20% heat-inactivated fetal bovine serum in Dulbecco phosphate-buffered saline plus 50 µg/ml penicillin/streptomycin, 50 µg/ml gentamicin and 2 µg/ml Fungizone [Sigma Aldrich, St. Louis, MO, USA]) containing a 4 mm bead (Daisy Rogers, Arkansas). Saliva was collected by inserting the proboscis of the female mosquitoes into a capillary tube containing ~20 µl fetal bovine serum plus 50% sucrose 1:1 for 30 minutes and subsequently ejecting the mixture into 125 µl MD. Mosquito bodies were then placed in individual tubes containing 500 µl MD and a bead. All samples were held at −80°C until assayed.

Infection, dissemination and transmission results were obtained by respectively screening the whole bodies, legs and saliva collected at different time points, as described by Ciota A.T et al., 2017. To obtain viral titer, ZIKV-specific quantitative PCR assay was utilized (19) as described in Ciota A.T et al., 2017.

### Statistical analysis

Statistical analysis was performed with GraphPad Prism version 5.0. Infection; dissemination and transmission rates were compared using Z tests, Fischer’s exact test and Chi-Square tests. The viral loads at the different stages of infection were compared using an ANOVA test. Generalized linear models (GLM) were used to determine the relative impact and interaction of different variables on vector competence.

### Vectorial capacity

Vectorial capacity which determines the intensity of pathogen transmission and incorporates blood feeding behavior and vector longevity along with vector competence was calculated in this study using below formula:

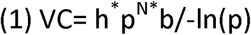

Where h = host feeding rate (proportions of mosquitoes acquiring at least two artificial blood meals in their life time), p = the probability of daily survival, p=(100-[-slope]) of the best-fit linear relationship between mortality and time to transmission. N = the mean extrinsic incubation period (EIP), calculated here as the mean days to dissemination and b = vector competence (proportion of exposed mosquitoes developing disseminated infections and capable of transmitting infection). The host feeding rate and the longevity were evaluated in a separate study using the same populations and temperature regimens. Briefly, six populations of *Aedes* mosquitoes were reared at similar temperature regimes and conditions as described above. The female mosquitoes were allowed to mate post adult emergence and after three days, placed in separate 3.8 L cartons and offered an uninfected blood meal. Blood meals were subsequently offered every three days nine times, until the last female mosquito died. An account was kept of mortality and feeding rates of female adult mosquitoes throughout the experiment. The linear regression analysis of these data was completed using GraphPad Prism 5.0.

## Results

### Vector competence

*Ae. aegypti* and *Ae. albopictus* were equally susceptible to ZIKV infection [Z-Score = 0.81 P(0.41794)]. Overrall, temperature influenced infection rates in a population and species-specific manner. Higher temperatures were associated with decrease ZIKV infection of AEG MI at both 4 and 14 dpi. AEG MI reared at the M temperature regimen had significantly higher proportions of individuals infected compared to the H temperature regimen at both 4 dpi (M = 0.9, H = 0.67, Z score = −2.1935, P = 0.02852) and 14 dpi (M = 1.0, H = 0.67, Z score = −3.4641, P = 0.00054). AEG MX, on the other hand, demonstrated no significant difference in infection rates between temperature regimes at 4-14 dpi. While there was also no significant effect of temperature on infection rates of ALB LI at 4 and 14 dpi, at 7 dpi ALB LI individuals reared at the M regime had a higher infection rate than the L regime (M = 0.96, L = 0.77, Z score = 2.2787, P = 0.0226; Figure 3).

**Figure 3.**
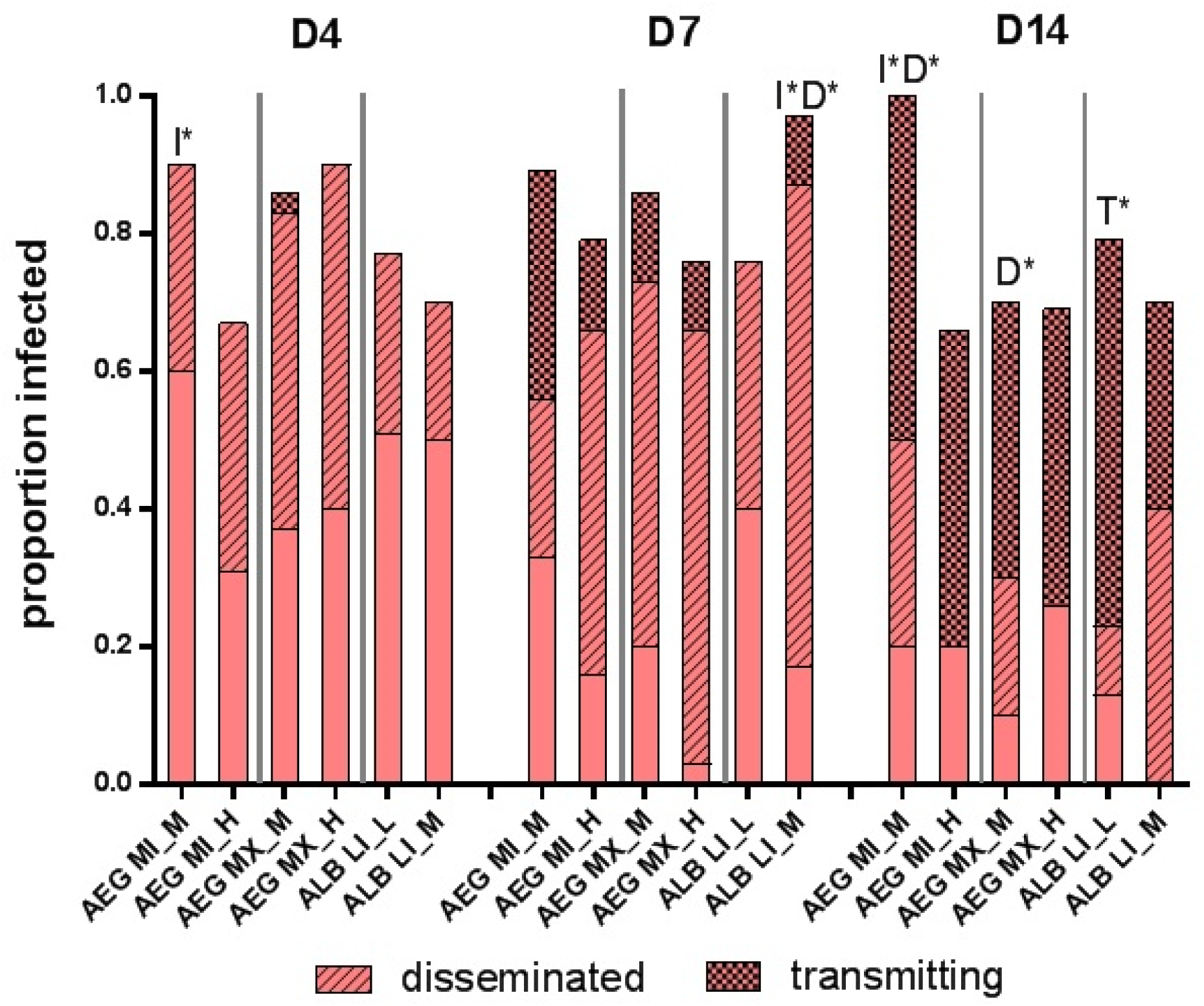
The infection and dissemination rates of *Aedes* mosquitoes infected with ZIKV and reared at different temperature regimes. The height of the bar plot represent the proportion of blood fed individuals successfully infected with ZIKA virus. The proportion of these that disseminated and subsequently transmitted the virus are further indicated. The AEG MI (M) infection rates were significantly higher than AEG MI (H) at both the earliest time point (4 dpi) and the last time point (14 dpi). At 7 dpi, ALB LI (M) infection and dissemination rates were significantly higher than ALB LI (L). The ALB LI (L) had the highest transmission efficiency.

Dissemination rates were influenced by population, temperature and extrinsic incubation period (Glm model P [0] Supplementary file no. 1). There was no significant differences noted in the dissemination rates except for the ALB LI at 7 dpi, (M = 0.83 and L = 0.45, Z-Score = 2.6671, P = 0.00758). The transmission efficiency of the ALB LI L population was highest at 14 dpi and was significantly different from the ALB LI M individuals (Z-Score = −2.0842, P = 0.03752). The other populations had similar transmission efficiencies among temperature regimes at all time points (Table 1; Figure 3).

**Table 1.** A table showing the infection, dissemination, transmission rates and transmission efficiency across populations, species and time points.

| Species | Geographical origin | Temp. regimen | % Infection rate(whole bodies ) | % Dissemination rate (no. of mosquitoes) | % Transmission rate (no. of mosquitoes) | %Transmission efficiency (no. of mosquitoes initially screened) |
| --- | --- | --- | --- | --- | --- | --- |
| **4 days post infection** |  |  |  |  |  |  |
| *Ae. aegypti* | Miami | D32N28 (H) | 67(30) | 55(20) | 0(11) | 0(30) |
| *Ae. aegypti* | Miami | D30N26 (M) | 90(30) | 37(27) | 0(10) | 0(30) |
| *Ae. aegypti* | Mexico | D32N28 (H) | 90(30) | 56(27) | 0(15) | 0(30) |
| *Ae. aegypti* | Mexico | D30N26 (M) | 90(30) | 56(27) | 13(15) | 7(30) |
| *Ae. albopictus* | Long Island | D30N26 (M) | 70(30) | 29(21) | 0(6) | 0(30) |
| *Ae. albopictus* | Long Island | D28N24 (L) | 77(30) | 35(23) | 0(8) | 0(30) |
| **7 days post infection** |  |  |  |  |  |  |
| *Ae. aegypti* | Miami | D32N28 (H) | 83(30) | 76(25) | 26(19) | 17(30) |
| *Ae. aegypti* | Miami | D30N26 (M) | 93(30) | 61(28) | 6(17) | 3(30) |
| *Ae. aegypti* | Mexico | D32N28 (H) | 77(30) | 96(23) | 14(22) | 10(30) |
| *Ae. aegypti* | Mexico | D30N26 (M) | 87(30) | 77(26) | 21(20) | 13(30) |
| *Ae. albopictus* | Long Island | D30N26 (M) | 97(30) | 86(29) | 12(25) | 10(30) |
| *Ae. albopictus* | Long Island | D28N24 (L) | 77(30) | 52(23) | 0(12) | 0(30) |
| **14 days post infection** |  |  |  |  |  |  |
| *Ae. aegypti* | Miami | D32N28 (H) | 67(30) | 80(15) | 78(18) | 47(30) |
| *Ae. aegypti* | Miami | D30N26 (M) | 100(30) | 83(30) | 60(25) | 50(30) |
| *Ae. aegypti* | Mexico | D32N28 (H) | 70(30) | 63(19) | 93(14) | 43(30) |
| *Ae. aegypti* | Mexico | D30N26 (M) | 73(30) | 86(22) | 68(19) | 43(30) |
| *Ae. albopictus* | Long Island | D30N26 (M) | 70(30) | 100(21) | 43(21) | 30(30) |

Interspecies differences in the mean viral load were measured at 4 and 14 dpi (Kruskal-Wallis test, P <0.0001). At 4 dpi, viral load in AEG MX (M) was significantly higher than ALB LI (L) (4.81 vs. 2.77 log_10_ pfu/ml). At 14 dpi, viral load had increased across all the populations. Notably, there was a significant effect of species on viral load (Kruskal-Wallis test P <0.00001), with viral load significantly higher in *Ae. aegypti* relative to *Ae. albopictus* across time points (Figure 4). No significant effect of population or temperature on viral load was measured.

**Figure 4.**
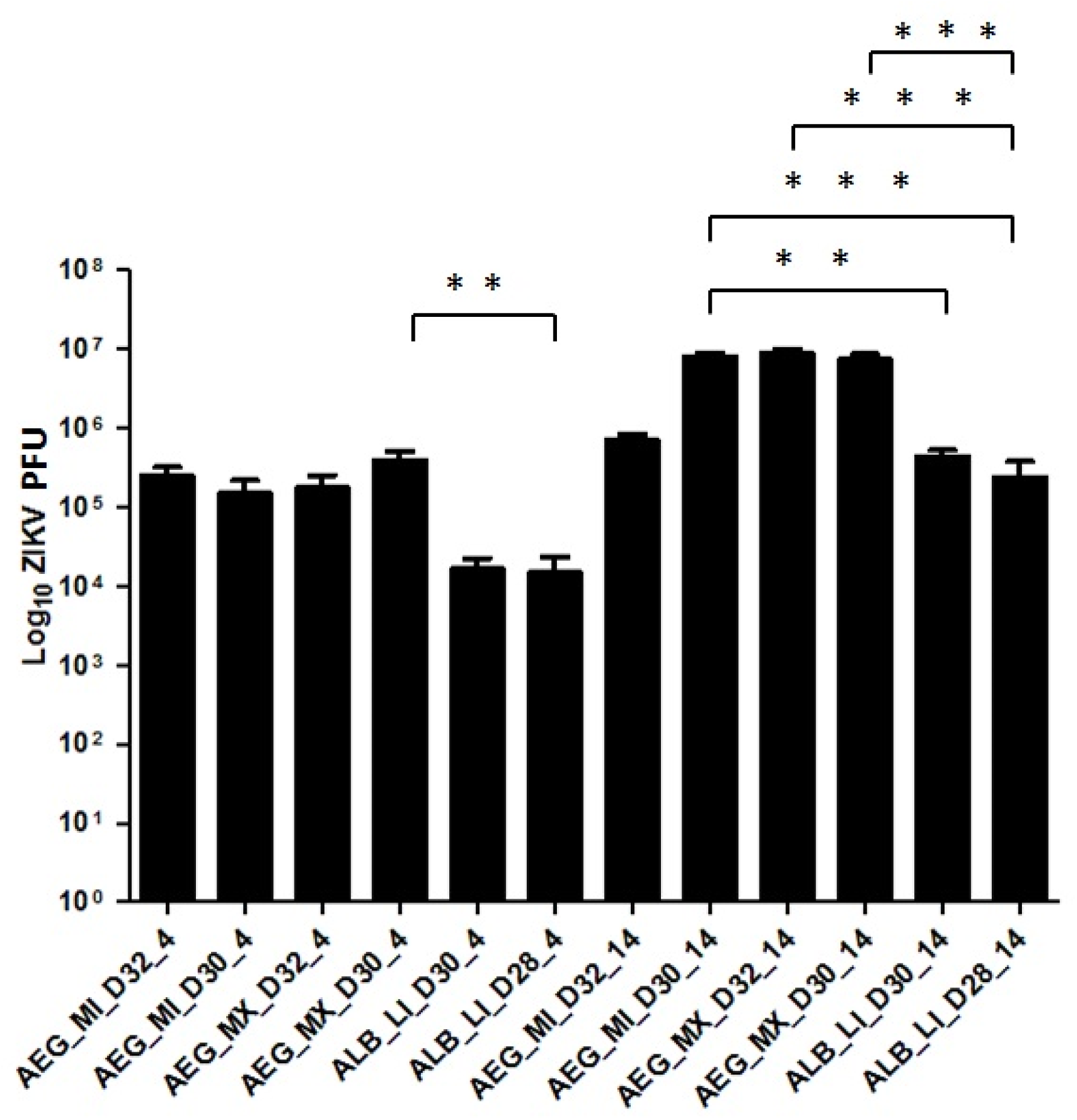
The absolute viral titer load comparison of ZIKV-infected *Aedes* mosquitoes at different temperature and time points. The plot represents the absolute viral titers of individuals of different populations of *Aedes* mosquitoes utilized in this study. The distribution of the viral titer around the mean is represented by the error bar.

### Vectorial capacity

Increase in temperature resulted in decreased frequency of blood feeding for both species of *Aedes* mosquitoes in this study (AEG [H(138/584)(0.23), M(209/732)(0.28)], Z score = −5.8478, P[0]; ALB [M(40/125)(0.32), L (162/303) (0.53), Z score = 4.0449, P[0]; (Figure 5). *Ae. aegypti* populations reared at the higher temperature regimens had a consistently reduced longevity relative to the low temperature regimen individuals (Log-rank [Mantel-Cox] test, Chi-square, df 35.66, 5, P <0.0001); Figure 6). *Ae. albopictus* had a significantly higher survivorship compared to both *Ae. aegypti* populations and longevity was not affected by temperature.

**Figure 5.**
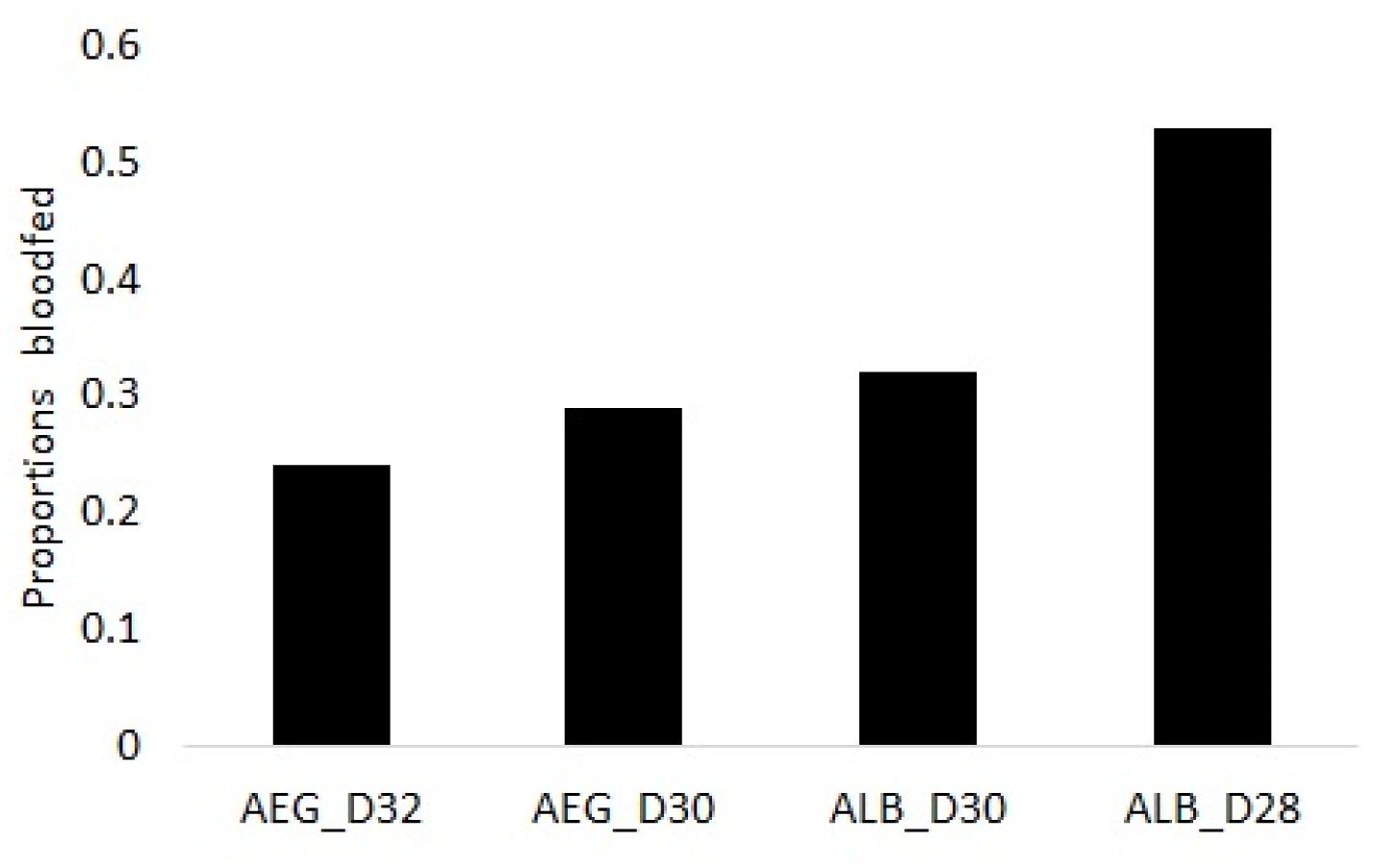
The proportion of blood fed individuals. The bar graphs represent female mosquito individuals reared in a separate experiment and blood fed with a non infectious blood meal post mating. The individuals were reared at similar temperature regimes as described above: AEG_D32 (AEG MI and AEG MX reared at H temperature regime); ALB_D30 (ALB LI reared at M temperature regime) while ALB_D28 (ALB LI reared at L temperature regime). There was significant heterogeneity in the number of blood fed individuals across the temperature regimes.

**Figure 6.**
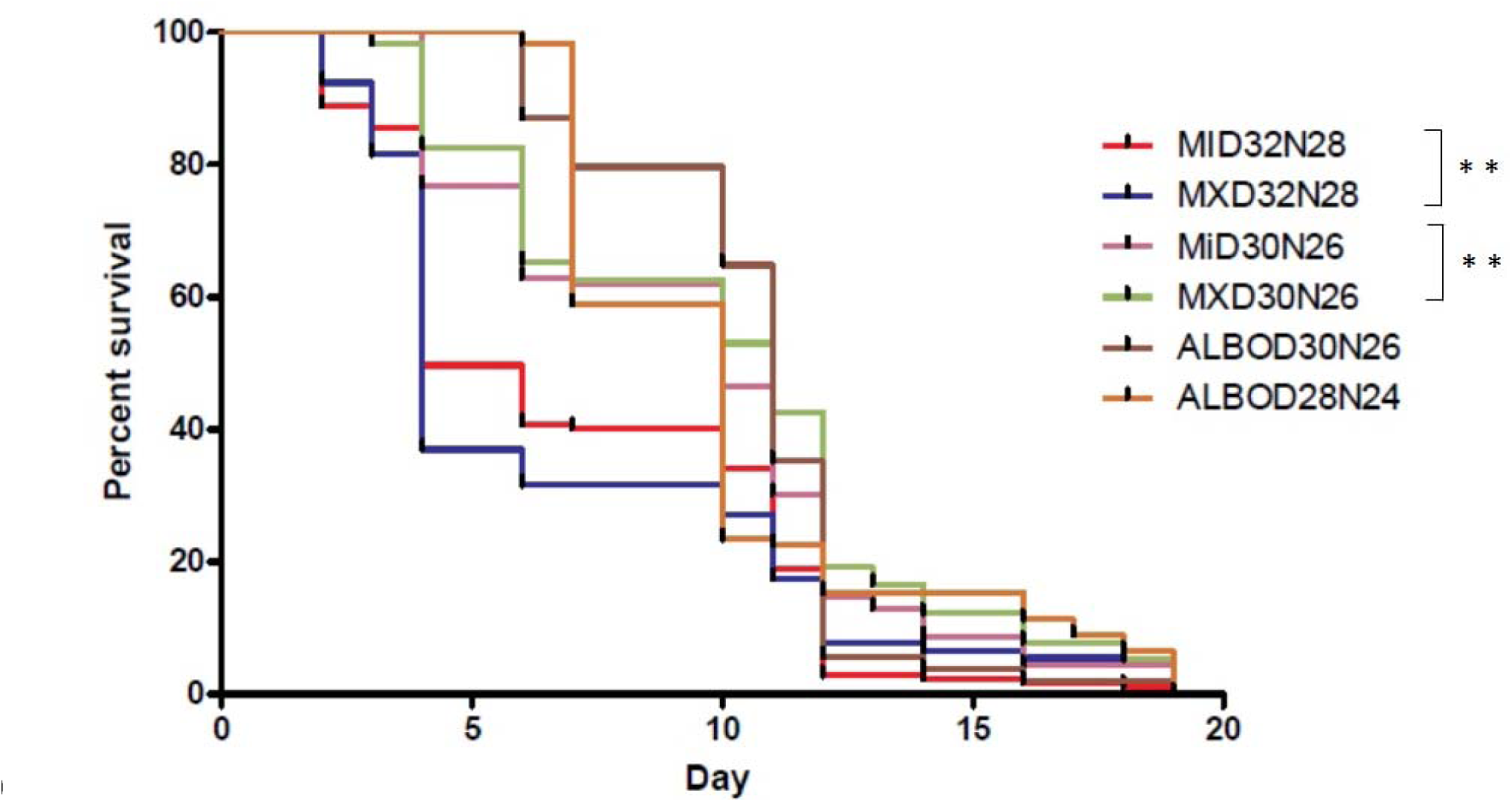
The survival curves of *Aedes* mosquitoes in response to rearing at varying temperature regimes. Female reared in a separate experiment under similar conditions and temperature regimen, blood fed and observed for survival rates post emergence. The Kaplan-Meier survival curves as measured days post emergence and analysed by the Logrank test revealed statistical significance between populations across the different temperature regimens.

Overall, ALB LI (L) had the highest vectorial capacity (1.63), while AEG MI(H) had the lowest (0.11). Across all populations, increasing temperatures decreased vectorial capacity (Table 2). It is noteworthy that the vectorial capacity of the *Ae. albopictus* individuals reared at M temperature regimen was higher than that of the *Ae. aegypti* also reared above the baseline peak transmission temperature ALB LI (M) = 0.64, AEG MI (H) = 0.11 AEG MX (H) = 0.16; Table 2). While this is driven largely by decreases in longevity and vector competence for *Ae. aegypti*, decreases in vectorial capacity of *Ae. albopictus* is primarily a result of decreased blood feeding at higher temperatures.

**Table 2.** Vectorial capacity of the *Aedes* mosquito populations for ZIKV.

| | Source | Temp. regimen | h | P | N | b | $VC=h^2P^Nb/-\ln(P)$ |
| --- | --- | --- | --- | --- | --- | --- | --- |
| AEG MI (H) | Miami | D32N28 | 0.21 | 0.92 | 8.21 | 0.4 | **0.11** |
| AEG MI (M) | Miami | D30N26 | 0.3 | 0.94 | 9.82 | 0.8 | **0.63** |
| AEG MX (H) | Mexico | D32N28 | 0.26 | 0.92 | 7.66 | 0.37 | **0.16** |
| AEG MX (M) | Mexico | D30N26 | 0.31 | 0.95 | 8.52 | 0.6 | **0.73** |
| ALB LI (M) | Long Island | D30N26 | 0.32 | 0.94 | 9.52 | 0.7 | **0.64** |
| ALB LI (L) | Long Island | D28N24 | 0.53 | 0.94 | 9.97 | 0.67 | **1.63** |

AEG- *Ae. aegypti*
ALB- *Ae. albopictus*
MI- Miami
MX- Mexico
LI- Long Island
h- host feeding rate
p- Probability of daily survival
N- Mean extrinsic incubation period
b- proportion of exposed mosquitoes developing disseminated infections

## Discussion

This study demonstrates that increasing temperatures could decrease vectorial capacity of *Aedes* mosquitoes for ZIKV. Measured decreases in vectorial capacity resulted from alterations to vector competence, blood feeding frequency and /or longevity. While the magnitude of these effects was dependent on species, population and temperature regime, an overall decrease in ZIKV transmissibility was measured with all three populations evaluated following a 2°C rise in diurnal temperature cycle. A larger decrease was measured for *Ae. aegypti* mosquitoes when modeling warmer current and predicted future temperature regimes of the southern US and Mexico. These results support the idea that future climate change could result in a northern shift in suitability of *Aedes* populations for transmission of ZIKV and other invasive arboviruses.

Higher temperatures decreased the blood feeding rates of the *Aedes* mosquitoes in this study. The mosquitoes in our study were provided with water and sugar *ad libitum* and both were suspended 24 h before offering a blood meal. There is a possibility that the mosquitoes reared at high temperature may have imbibed excess water and sugar to avoid dehydration and this could have in turn led to decreased intake of blood (or less likelihood to feed on blood). Mosquitoes that had access to sugar before blood meals demonstrated deferred blood meal intake compared to the sugar deprived (20). In addition, *Anopheles gambiae* and *Ae. aegypti* have been shown to forego nectar feeding and instead imbibe multiple blood meals during each gonotrophic cycle (21–23). A caveat to our results is that in our study, we offered the mosquitoes sweetened defribinated blood, hence *in vivo* blood meal might yield different results. Future studies more precisely monitoring feeding behavior could clarify this yet, regardless of mechanism. These results suggest that climate change may have a significant effect on transmission risk of *Aedes* mosquitoes by altering their frequency of blood meal acquisition. The decreased longevity due to temperature measured in this study may have resulted from the negative impact that higher temperature had on mosquito physiology (14). Increase in body temperature of an insect results in increase of both metabolism and respiration upto a critical thermal limit. Reports have demonstrated that death occurs soon after respiration rate begin to drop, in spite of the insect being returned to normal temperatures, this may be indicative of systemic cell deaths at the high temperatures (24). The increased temperature may also negatively impact the nervous and endocrine systems controlling the insect metarmorphosis (24) which could subsequently lead to developmental defects leading to death (25).

Previous studies by and large demonstrate that shorter EIPs and overall increases in competence are associated with increases in temperature (2,26,27). Although our data generally support the idea that increasing temperatures accelerate EIP, overall competence was either unchanged (ALB LI and AEG MX) or decreased (AEG MI). While this is to some extent population-dependent, it may also be attributed to the use of fluctuating rather than constant temperatures ^29,^(29). In addition, immature stages also were subject to fluctuating temperature regimes, which could potentially have altered gene expression prior to exposure. The effect of temperature on infection rates was population and species-specific such that the AEG MI population had higher overall infection rates at the lower (M) temperature regimen, while there was no difference in infection rates of the AEG MX population between temperature regimes at any time point. *Ae. albopictus*, on the other hand, had higher infection and dissemination rates at 7 dpi when held at higher temperatures. While this difference is of interest, it is important to note that the higher temperature regime for *Ae. albopictus* (M) was equivalent to the baseline temperature of *Ae. aegypti*. It is unclear if further temperature increases would result in similar decreases of competence as were measured in AEG MI. Our results are comparable to Tesla et al., 2018 which demonstrated that ZIKV transmission by *Ae. aegypti* was optimized at a mean temperature of approximately 29°C when mosquitoes were held at constant temperatures ranging from 16°C-38°C. Taken together, these data demonstrate that temperature is a significant component of ZIKV transmission; and defining optimum conditions for individual populations and species is important for an accurate prediction of how future climate change will affect spatial expansion and transmission. Viral loads in *Ae. albopictus* were significantly lower than *Ae. aegypti* across time points. Our results corroborate Ciota et al., 2017 findings conducted at 28°C. Previous studies have demonstrated *Ae. albopictus* is a competent vector for ZIKV in the laboratory (1,8). However, the vector competence has been shown to be influenced by the ZIKV strain and the spatial origin of the mosquito population (1).

The explosive spread of ZIKV in the Americas raised concerns that another vector in addition to *Ae. aegypti* might be involved in ZIKV transmission(31). Our study confirms the potential for *Ae. albopictus* to play a role in ZIKV epidemics in the Americas, although they have not been demonstrated to be significantly involved to date.

However, since variation in vector competence among *Ae. aegypti* populations and virus strains has been reported previously (8,28,32,33), and our results further support this, a comprehensive assessment of the potential effect of climate change would require additional studies with multiple populations and ZIKV strains.

There are two important caveats to this study. First, it is unclear how mosquitoes will adapt or evolve in response to incremental changes in temperature and this could significantly alter the vectorial capacity of future popualtions. Second, alterations to life-history traits including blood feeding behavior and longevity can result from arbovirus infection (34).

We measured lower infection rates at later timepoints, particularly at high temperatures. Since arboviral infections in mosquitoes are persistent and generally are not cleared, these data suggest that ZIKV infected indivduals were less likely to survive the experimental period and that this increased mortality was facilitated by increased temperatures. Further studies monitoring life-history traits following infection at altered temperatures will help clarify this relationship.

Despite these caveats, the results of our study suggest that future global climate change resulting in increased global temperature will have negative fitness cost on *Aedes aegypti* mosquitoes for ZIKV and could significantly alter their vector competence. Further, the impact of future climate change will be driven by unique intrinsic interractions between specific vectors and pathogens and therefore is unlikely to be uniform across different species and populations.(30)

## Supporting information

https://figshare.com/account/home

## Acknowledgements

This publication was supported by the Cooperative Agreement Number U01CK000509 funded by the Centres for Disease Control and Prevention. Its content are solely the responsibility of the authors and do not necessarily represent the official views of the Centres for Disease Control and Prevention or the Department of Health and Human Services. We would like to acknowledge the insightful discussions and comments by Dr. Constentine Dieme.

## Contributions

MGO: Involved in the study design, conducted the experiments, analyzed the results and drafted the paper

BS: Involved in the mosquito rearing, assisted in mosquito manipulation exercises

PA: Assisted in mosquito manipulation exercises

MN: Involved in the mosquito rearing, assisted in mosquito manipulation exercises

KL: Generated and propagated the ZIKV infectious clone utilised in this study

AV: Designed the linear regression model

CAT: Involved in the study design, participated in the writing of the paper

KLD: Involved in the study design, participated in the writing of the paper

## Supplementary file

**Supplementary file no. 1 (Available on** https://figshare.com/account/home). A GLM model showing the interactions between different variables and their effect on transmission rate, transmission efficiency, infection rate and dissemination rates of the *Aedes* mosquito populations reared at different temperature regimen and collected from different geographical origins. The significance of their effect was tested using ANOVA test.

